# Visual cortical dynamics supporting predictable attentional capture

**DOI:** 10.64898/2026.04.23.720330

**Authors:** Jisk J. Groot, Jeffrey D. Schall, Jacob A. Westerberg

**Affiliations:** Department of Vision and Cognition, Netherlands Institute for Neuroscience, Royal Netherlands Academy of Arts and Sciences, Amsterdam, Netherlands; Institute for Interdisciplinary Studies, University of Amsterdam, Amsterdam, Netherlands; Centre for Vision Research, Centre for Integrative and Applied Neuroscience, Department of Biology, York University, Toronto, Canada; Department of Psychology, Center for Integrative and Cognitive Neuroscience, Vanderbilt Brain Institute, Vanderbilt Vision Research Center, Vanderbilt University, Nashville, United States

**Keywords:** Attention, Distractor suppression, Macaque monkey, Population spiking, Predictive processing, Priming, Visual search

## Abstract

Visual behavior depends on the ability to prioritize relevant sensory information while filtering out distractions. Predictable sensory contexts enable more efficient behavior by altering sensory processing. Using laminar neurophysiology in macaque visual cortex measuring population spiking during a feature-based pop-out visual search task, we examined how predictable visual routines influence cortical columnar processing of sensory information. By manipulating predictability through attentional priming, we found improved behavioral performance with predictable stimulus arrays. These behavioral changes were supported by earlier neuronal attentional target selection driven by reduced variability in the sensory response to the predictable target stimulus and more homogeneous feedforward processing dynamics across the cortical column. More effective distractor suppression was driven by adapted feedforward processing of the more frequent distractor stimulus. Together, these changes implicate independent mechanisms for target enhancement and distractor suppression. Our results highlight how predictions from experience alter visual columnar processing to optimize attentional selection through streamlining feedforward signaling.

**Highlights:**

- Both target enhancement and distractor suppression are independently observable in visual cortex with different profiles of modulation as a function of predictability.
- Predictions produce stronger, more reliable feedforward responses to attention-capturing targets across the visual cortical layers.
- Better distractor suppression with stimulus predictability is driven by adapted sensory representations, not top-down inhibition.

**Significance:** How do predictions help determine what catches our attention? It is well known that when primates repeatedly perform visuomotor tasks, they become more proficient. Using neural recordings across the layers of the visual cortex, Groot et al. demonstrate that more efficient feedforward processing of sensory information contributes to improved visual behavior under predictable sensory conditions. These findings highlight the crucial role of predictions in proactively shaping sensory representations to facilitate efficient attentive behavior.

## Introduction

Our senses continuously sample a broad range of rich sensory information. Yet, the brain’s limited processing capacity prevents it from scrutinizing every input. As such, efficient interaction with the environment requires prioritizing behaviorally relevant information while filtering out irrelevant inputs. Neuronal operations mediate this selective allocation of processing resources, generally referred to as attention [1–32]. However, attention benefits from, and likely gates [33], another operation, prediction. Predictions about the sensory context can guide attention to stimuli expected to be more valuable [34–37]. Thus, the interaction between attention and prediction forms a critical foundation for efficiently distilling rich sensory input into actionable information.

One form of attention, stimulus-driven attention [38], is characterized by salient stimulus features that capture attention [39–41]—for instance, a brightly colored road sign or a notification sound from your phone. Attentional capture is supported by feedforward processing of sensory information such that sensory representations containing salient information are enhanced relative to competing sensory representations [42]. However, attentional capture is dynamic; tasks that benefit from it, such as pop-out visual search, notably benefit from predictable stimulus sequences. This is evident in a special case of pop-out visual search, known as priming of pop-out (PoP), in which behavioral performance improves with repetitive stimulus sequences. The pop-out stimulus feature, which engenders attentional capture, becomes predictable with repetition, and reaction times fall [43–45]. This form of prediction does not dictate the arrangement of sensory stimuli, but rather represents a more general, predictable sensory context. In other words, predictions are made about which stimuli to expect, but their precise locations are unpredictable. However, it is not clear to what extent changes in sensory processing in predictable sensory contexts drive behavioral changes, or whether the associated behavioral improvements result entirely from changes in processing downstream of early sensory representations [46].

Earlier work has implicated sensory areas in prediction-enabled improvements in attentional capture tasks [42,47– 50]. Notably, loss of sensory representations selective for the attention-capturing feature damages the behavioral benefits associated with stimulus predictability [47]. Moreover, predictable stimulus sequences induced changes in sensory representations that were selective for the salient feature, promoting them for attentional capture. Critically, these changes were already measurable before the stimuli appeared [42]. As such, the question is not whether the sensory cortex contributes, but rather how predictions reshape sensory processing to subsequently achieve more efficient attentive behavior. In predictable visual search conditions, a crucial question is whether changes in sensory processing are already present in the bottom-up response, or whether behavioral improvements follow the more effective top-down deployment of spatially selective attention to the target stimulus. One avenue to interrogate this is through exploiting the functional neuroanatomy of cortical columnar microcircuits [51,52].

Neuronal processing in sensory areas leverages the laminar structure of cortical tissue to enable mesoscopic, circuit-level computations [35,51–54]. Importantly, bottom-up versus top-down processes can be tracked across cortical columns, exhibiting predictable and dissociable temporal and spatial signatures [18,21,26,36,55,56]. We sought to investigate how a predictable sensory context changes neural processing along cortical columns in the visual cortex, thereby promoting more effective attentional capture. Specifically, we tasked macaque monkeys with performing a color-based priming of pop-out task while neural activity was recorded across the layers of mid-level visual cortical area V4. We contextualize priming of pop-out as a predictive-processing task in which stimulus sequences become predictable with repetition, thereby forming a predictable sensory context. When the attention-capturing features change, we defy those predictions. We then tested whether predictability primarily affects the processing of distracting information or the attended information to improve behavior. We examine epochs of predominantly feedforward vs. recurrent processing to determine whether changes are already present in feedforward processing or arise from putative feedback. We find that it is primarily more streamlined feedforward processing of sensory information, rather than more effective top-down selective attentional enhancement or distractor suppression, that subserves predictable-context-enabled attentive behavior.

## Results

### Task and behavior

To investigate how sensory processing changes with predictable sensory contexts during attentive behavior, multiunit activity (MUA) was recorded across the cortical layers of area V4 in two macaque monkeys (Ca and He) performing a color-based priming of pop-out visual search task (Figure 1A). On each trial, the monkeys made an eye movement to an oddball stimulus (target) defined by its color (red or green) among five uniform distractors of the opposite color. Importantly, the color identity of the target and distractors remained constant within blocks of consecutive trials, with the target-defining feature switching unpredictably between blocks (Figure 1B).

**Figure 1:**
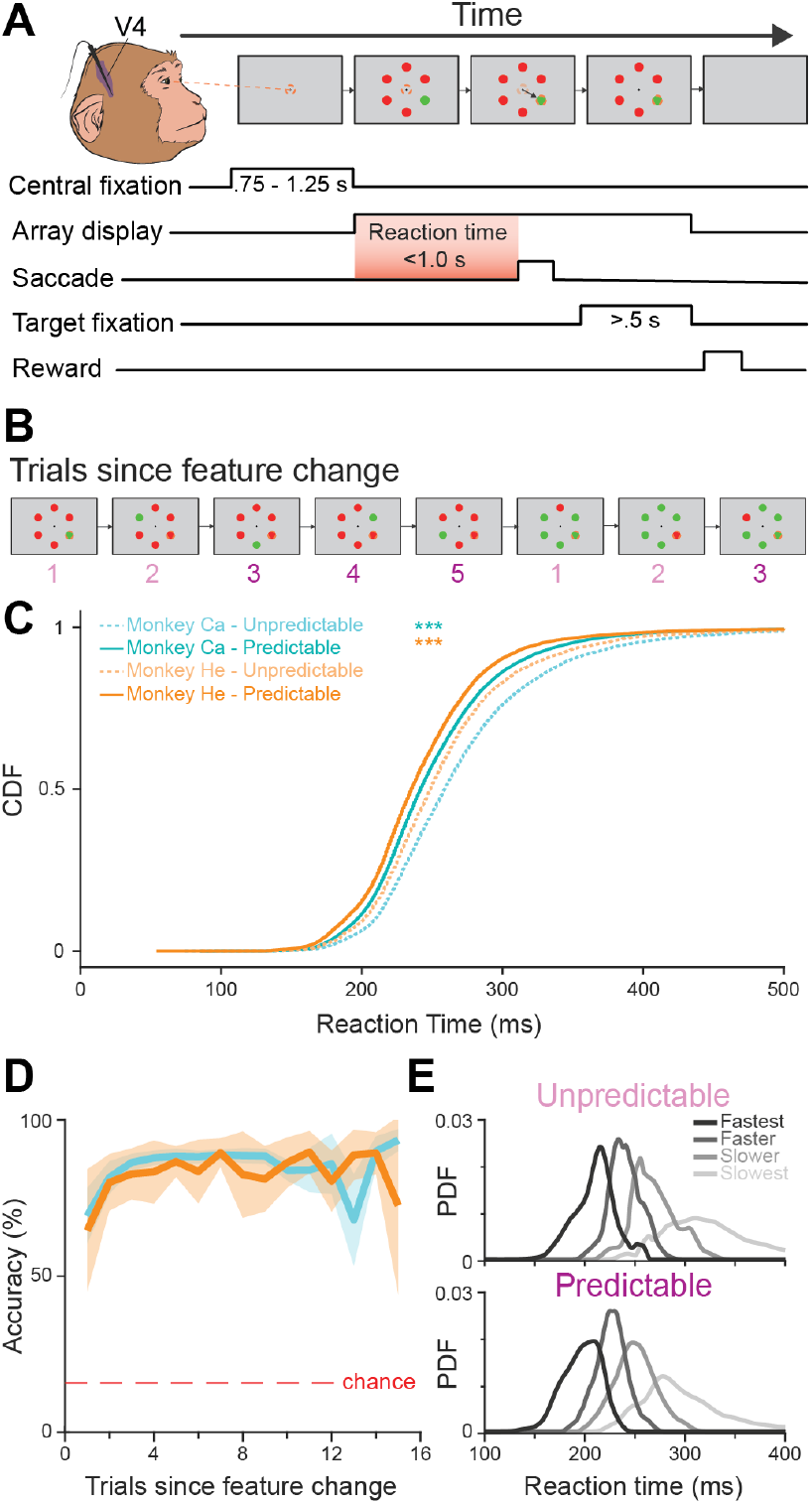
Behavioral task and performance. **A)** Monkeys fixate on a central cross for a variable period, then an array of 6 colored disks appears. 5 disks were distractors of the same color, and 1 was a uniquely colored “oddball” (target). Monkeys made a saccade toward the target disk and held fixation for 500 ms. Correct choices were rewarded. **B)** Target and distractor colors remained fixed within a block of trials, then swapped to begin a new block. Trials 1-2 following a color change were defined as “unpredictable”; trials 3-15 were classified as “predictable”. **C)** Reaction time distributions: Cumulative distribution functions (CDFs) of reaction times for each monkey (Ca: turquoise, He: orange), separated by unpredictable (trials 1-2) and predictable (trials 3-15) contexts. Differences in RT distributions between unpredictable and predictable contexts were tested using the Wilcoxon signed-rank test (α = 0.05). **D)** Task accuracy plotted as a function of trial number since the color change for both monkeys (Ca: turquoise; He: orange). 95% confidence intervals surround each mean. The dotted red line indicates the chance level (16.7%). **E)** Reaction time distributions: Probability distribution functions (PDFs) of reaction times, segmented into quartiles, are shown for unpredictable and predictable contexts.

Data were collected across 29 recording sessions (Ca: n = 21; He: n = 8), and analyses were limited to trials where the monkey correctly identified the target. We focused on the first 15 trials after each feature change. “Unpredictable” trials were defined as the first two trials after a change in the target color [48,57,58], where the target feature was relatively new, and the oddball-defining feature was not yet, or only minimally, predictable. The “Predictable” condition included trials 3-15, where the target-defining feature had been repeated more than once within the current block.

Both monkeys showed strong behavioral facilitation once the predictable context was established: Response times were significantly shorter in the predictable context compared to the unpredictable one (Ca: median = 241.99 ms, 95% CI = [241.05, 242.03]; unpredictable = 258.99 ms, 95% CI = [258.01, 260.01]; rank sum z = 37.94, p = 5.41 × 10^−315^. He: median = 235.03 ms, 95% CI = [234.05, 236.01]; unpredictable = 248.99 ms, 95% CI = [247.03, 250.02]; rank sum z = 16.03, p = 8.42 × 10^−58^), indicating faster deployment of attention (Figure 1C). Furthermore, accuracy increased significantly after the first repeated trial (Figure 1D), as confirmed by a repeated-measures ANOVA (α = 0.05), which showed a significant main effect of trial on performance (F(14, 28) = 2116.2, p = 1.11 × 10^−16^). Post-hoc comparisons revealed that accuracy in the second trials was significantly higher than in the first trials (p = 0.0102), with an average improvement of 19.27% (± 3.58% SEM). This facilitation continued with further repetitions, with significant increases between the 1^st^ and 3^rd^-9^th^ trials (p < 0.02 for each), before reaching a plateau from the 10^th^ to 15^th^ trials (p > 0.05).

As prior work indicated that stratifying neural data into behavior-associated bins uncovers hidden neural dynamics [42], we quartiled reaction times per session, yielding four behavioral groups per predictive context (Figure 1E). This grouping enables a systematic comparison of behavioral and neural dynamics across the range of observed response times, helping to identify patterns of neural activity that could underlie distinct behavioral outcomes. We return to these behavior-associated bins in later analyses.

### Laminar dynamics in (un)predictable search conditions

At the neuronal level, attentional selection influences sensory processing through two complementary processes: target enhancement (TE), in which responses to behaviorally relevant stimuli are increased, and distractor suppression (DS), in which responses to irrelevant stimuli are decreased [42,46,48,59,60]. Importantly, these modulations improve neural discriminability between targets and distractors, leading to faster target selection times (TST), i.e., the instant at which neural activity differentiates behaviorally relevant and irrelevant stimuli [2]. These TSTs decrease with predictability during the priming of the pop-out (PoP) task we employ [46,48,57]. However, the source of these reduced TSTs in this task remains unknown. We first examine TSTs across cortical layers to identify the origins of these changes.

To determine how the laminar circuitry of target selection in visual cortex V4 contributes to behavioral performance and how these dynamics are modulated as sensory context becomes predictable, we analyzed neural activity across cortical layers during attentional selection under both the unpredictable and predictable contexts of the PoP task (Figure 2). Linear multielectrode arrays were positioned in V4, and putative laminar compartments were labeled using current source density (CSD) analysis, allowing electrodes to be grouped into upper, middle, and deep compartments [62–65]. Session-by-session assignment of data to the laminar compartment and alignment of data relative to the granular-infragranular boundary were reported previously [21,66]. Target selection was estimated as the time, relative to array onset, at which the neural population encoding the target stimulus showed significantly higher neuronal activation than the neural population encoding the distractor stimuli (running rank-sum test, α = 0.01, minimum cluster length = 25 ms). During both target and distractor presentations, target selection emerged in all layers under both unpredictable and predictable contexts (Figure 2; “All” column). In the unpredictable context, TSTs were comparable across layers (upper: 138.2 ± 2.77 ms; middle: 133.8 ± 4.76 ms; deep: 133.4 ± 8.20 ms). In the predictable context, TSTs were markedly reduced (upper: 84.6 ± 6.62 ms; middle: 82.0 ± 5.61 ms; deep: 86.6 ± 7.40 ms). Two-sided Wilcoxon signed-rank tests (α = 0.01) confirmed significant reductions in the upper (W = 449.0, p = 0.00960), middle (W = 1559.0, p = 2.13×10^−6^), and deep layers (W = 691.5, p = 7.19 × 10^−4^). This indicates that once the predictable context is established, target selection occurs earlier across all cortical layers.

**Figure 2.**
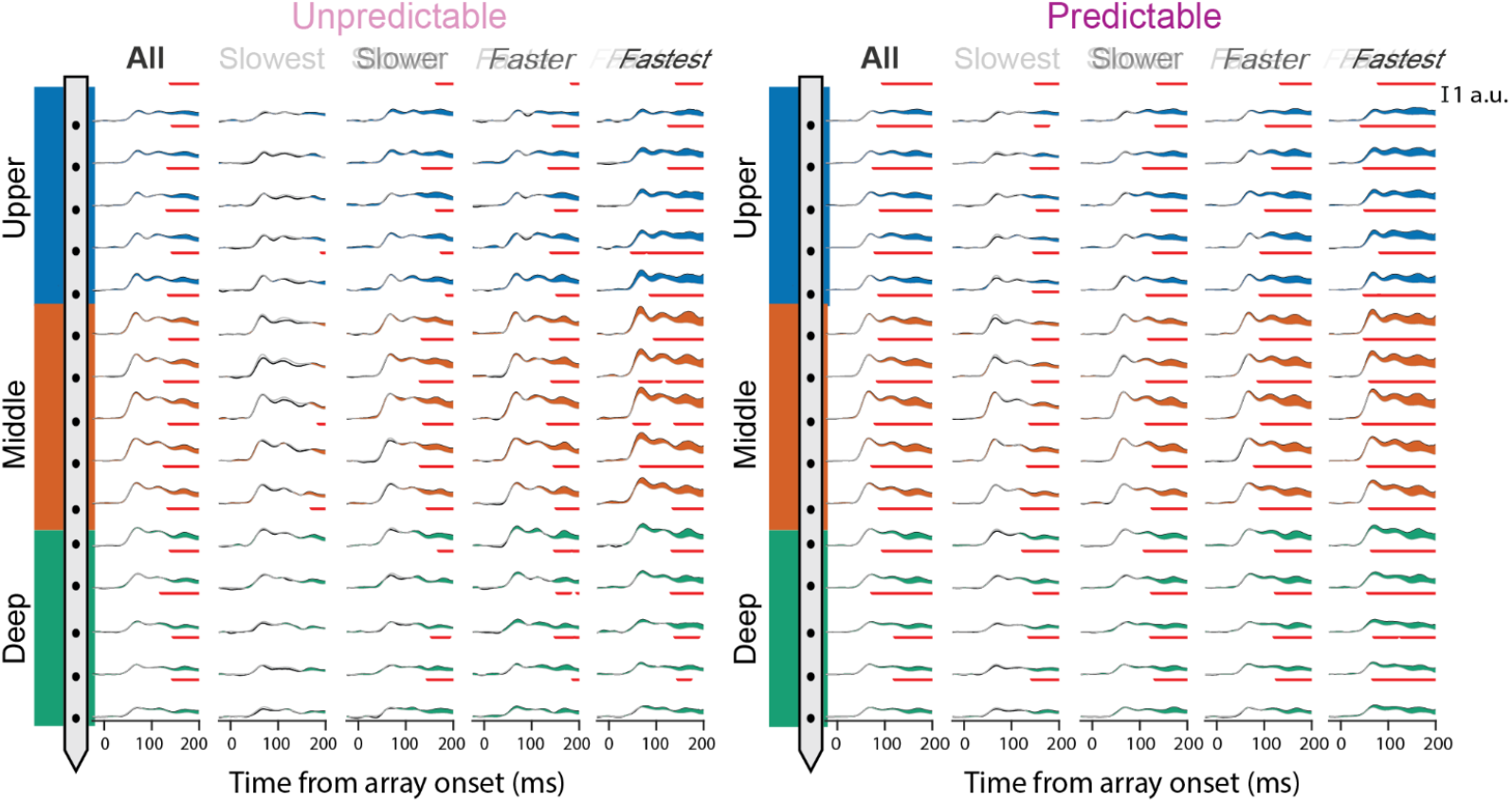
Neuronal attentional selection dynamics across depth and predictability context. Average multiunit activity (MUA) recorded for ™25 to +200 ms around stimulus array onset is shown for each channel, grouped by cortical layer (upper: blue, middle: orange, deep: green). Data presented separately for unpredictable (left) and predictable (right) contexts. For each context, responses are further divided by reaction time quartile, ranging from slowest (left columns) to fastest (right columns) trials, as well as all trials combined (“All”, first column). Within each panel, black traces indicate target responses, gray traces show distractor responses, and colored regions highlight the difference between target and distractor responses. Red bars indicate significant differences between target and distractor traces estimated by a running Wilcoxon rank sum test (α = 0.01, minimum cluster length = 25 ms). Scale bar at top right.

To further examine whether differences in TSTs were related to behavioral outcome, we compared TST profiles for trials grouped by reaction time (Figure 2; “Slowest” to “Fastest” columns). This revealed a clear pattern: faster reaction-time trials showed earlier target selection than slower trials in both the unpredictable and predictable conditions (see Table 1 for average target selection times across laminar compartments, stratified by behavioral performance). This reinforces the notion that rapid sensory target selection underlies fast reaction times. In addition, we observe a strong feedforward processing signature in the fastest trials during the unpredictable context, with selection emerging in the transient response first in the middle layer and then spreading to upper and deep layers. When the sensory context became predictable, target selection occurred more robustly across all laminar compartments. Layer-averaged target selection profiles further elucidated the underlying dynamics of target selection (Figure 3A).

**Table 1.** Average target selection times across cortical layers and predictability contexts. Average ± standard deviation of target selection onset (ms) across the three laminar compartments and stratified by behavioral performance (“Slowest” – “Fastest”). Dashes (™) indicate only one channel reached significance. UP = Unpredictable context; P = Predictable context.

| Layer | All |  | Slowest |  | Slower |  | Faster |  | Fastest |  |
| --- | --- | --- | --- | --- | --- | --- | --- | --- | --- | --- |
|  | UP | P | UP | P | UP | P | UP | P | UP | P |
| Upper | 140.6 ± 4.0 | 87.2 ± 7.1 | 194.0 ± – | 150.4 ± 4.7 | 167.4 ± 18.0 | 128.8 ± 11.4 | 153.2 ± 18.2 | 112.6 ± 15.7 | 115.6 ± 37.7 | 67.5 ± 17.5 |
| Middle | 134.6 ± 4.3 | 84.4 ± 5.8 | 179.5 ± 10.6 | 142.3 ± 5.1 | 137.8 ± 5.4 | 120.0 ± 5.7 | 145.0 ± 7.0 | 90.8 ± 5.3 | 71.6 ± 15.7 | 54.8 ± 6.1 |
| Deep | 138.8 ± 11.5 | 101.4 ± 20.5 | 172.0 ± – | 136.2 ± 10.9 | 158.0 ± 12.1 | 125.0 ± 8.2 | 159.6 ± 16.2 | 119.2 ± 14.9 | 139.2 ± 4.8 | 66.2 ± 5.0 |

**Figure 3.**
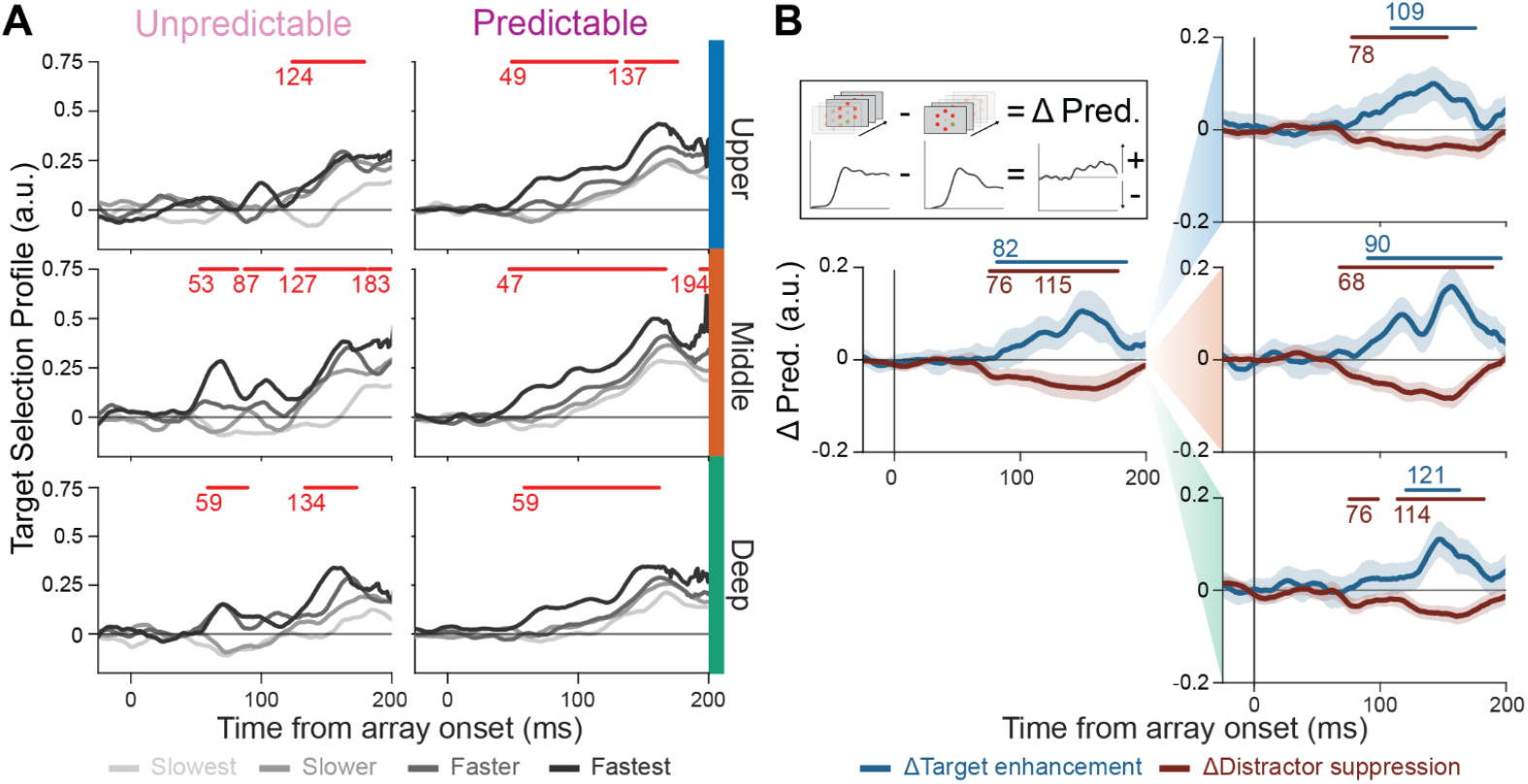
Laminar target selection dynamics become more homogeneous with predictability. **A)** Target selection profiles in unpredictable (left columns) and predictable (right columns) conditions, estimated by subtracting the distractor response from the target response for each trial. Profiles are shown separately by reaction time quartile (“behavioral outcome”; slowest to fastest) and averaged within each cortical layer across sessions. Red bars denote periods where traces corresponding to different behavioral outcomes (i.e., RT quartile) diverge significantly, as determined by a running Kruskal-Wallis test (α = 0.01, minimum cluster length = 25 ms). **B)** Changes in target enhancement (ΔTE: blue) and distractor suppression (ΔDS: red) profiles across laminar compartments. ΔTE and ΔDS are estimated by subtracting the average response during the unpredictable context from the predictable context. The left column displays the data averaged over all trials, the right column displays the data separated by laminar depth (top: upper; middle: middle; bottom: deep). Significance bars denote periods where traces corresponding to ΔTE and ΔDS diverge significantly from baseline (™250 to 0 ms before visual array onset), as determined by a running rank-sum test (α = 0.01, minimum cluster length = 20 ms). The inset illustrates how ΔTE and ΔDS are computed.

In the middle and deep layers, the fastest unpredictable trials displayed biphasic target selection profiles– exhibiting a first peak during the initial visual onset period (∼50-60 ms), followed by a second peak approximately 130 ms after visual array onset. This pattern suggests that both robust feedforward input (the early peak) and later recurrent enhancement (the later peak) contributed to the rapid selection in these trials. Interestingly, that early selection is not present in the upper layers, indicating that the selection signal may not be reliably fed forward through the columnar microcircuit, perhaps a reason for slower RT on those unpredictable trials [35].

Notably, the slowest trials in the unpredictable context showed an initial reduction in the target selection profile, with distractor representations briefly dominating the target. This pattern is consistent with a possible misallocation of attention in the feedforward window, perhaps another stimulus erroneously capturing attention. This phenomenon may be explained by negative priming, in which a previously ignored distractor feature becomes the target on subsequent trials, leading to reliably slower responses [67].

By contrast, in the predictable context, target selection profiles displayed earlier and more homogenous responses across all three laminar compartments. The upper layers showed an early increase in target selection, closely matching the early, more synchronous TST onsets across layers, which was notably absent in the upper compartment in the unpredictable context. This pattern was especially apparent in fast-response trials, in which target selection in the upper compartment occurred concurrently with that in the middle and deep compartments, creating a more uniform columnar response when stimulus identity was predictable. The greater columnar homogeneity and the earlier onset of target selection under predictable conditions suggest that predictability modulates local processing dynamics, potentially facilitating a more streamlined feedforward propagation of responses across cortical layers to support useful attentional capture.

### Target enhancement and distractor suppression across laminar compartments

Target selection is the effective difference between target and distractor responses, so changes in target selection can be mediated by either change in TE or DS. We refer to the changes in TE and DS as ΔTE and ΔDS. To investigate the ΔTE and ΔDS profiles, we subtracted the average response profiles for unpredictable contexts from those for predictable contexts, yielding independent contrasts for target and distractor trials (Figure 3B). The general cortical profile (left) shows clear ΔTE and ΔDS effects, with ΔDS (running rank-sum test, p < 0.01; mean onset time = 76 ms) initiating slightly before ΔTE (running rank-sum test, p < 0.01; mean onset time = 82 ms). This analysis reveals two key findings: (1) The different dynamics of ΔTE and ΔDS indicate that target enhancement and distractor suppression are independent mechanisms, and (2) The concurrent presence of both ΔTE and ΔDS provides evidence that predictability engages both mechanisms simultaneously, rather than selectively recruiting one.

Separation by laminar compartment (right) shows ΔTE and ΔDS across all layers, with the strongest effects in the middle layer. ΔTE seems to occur first in the middle (running rank-sum test, p < 0.01; mean onset time = 90 ms), followed by a slightly delayed onset in the upper layers (running rank-sum test, p < 0.01; mean onset time = 109 ms) and deep layers (running rank-sum test, p < 0.01; mean onset time = 121 ms).. However, it is worth noting that ΔTE appears in the upper layers around the same time as in the other two compartments but does not reach significance until later. ΔDS displays a relatively uniform onset of differences across laminar compartments, with the putative feedforward recipient middle layers showing slightly earlier ΔDS than the upper and deep compartments (running rank-sum test, p < 0.01; mean onset time: upper = 78 ms, middle = 70 ms, deep = 76 ms).

### Distractor suppression via neural adaptation

The early onset of ΔDS across all laminar compartments indicates that distractor responses are suppressed by changes in feedforward information processing rather than by active suppression via top-down modulation [68]. One possible underlying mechanism is neural adaptation [69]. Since the target is defined by a singleton feature among five homogeneous distractors per trial, and that structure is preserved across a block, the receptive fields (RFs) of neurons are more frequently exposed to the distractor feature within that block. This repeated exposure to the same distractor may lead to neural adaptation of that feature and therefore engender lower population responses across the block as the context becomes more predictable. This adaptation-induced response reduction could make up a portion of the change in distractor suppression we observe.

To test whether the ΔDS we observed reflects neural adaptation, we analyzed sequences of trials with consecutive distractor presentations and compared neural responses at each presentation (Figure 4). Specifically, we identified trial sequences that began with a target stimulus in the RF during a predictable context, followed by four consecutive distractor trials within the same target-feature block (Figure 4A). We included sessions with at least 15 occurrences, resulting in a total of 289 occurrences across 8 sessions (Ca, n = 196 across 6 sessions; He, n = 93 across 2 sessions). Using this selection, we compared the average response profiles across successive distractor-stimulus presentations to our neural population RF (Figure 4C). We observed a gradual reduction in neural response with distractor repetition, in both the middle and deep layers of cortex (Figure 4C). Notably, this pattern was already visible during the initial feedforward period.

**Figure 4.**
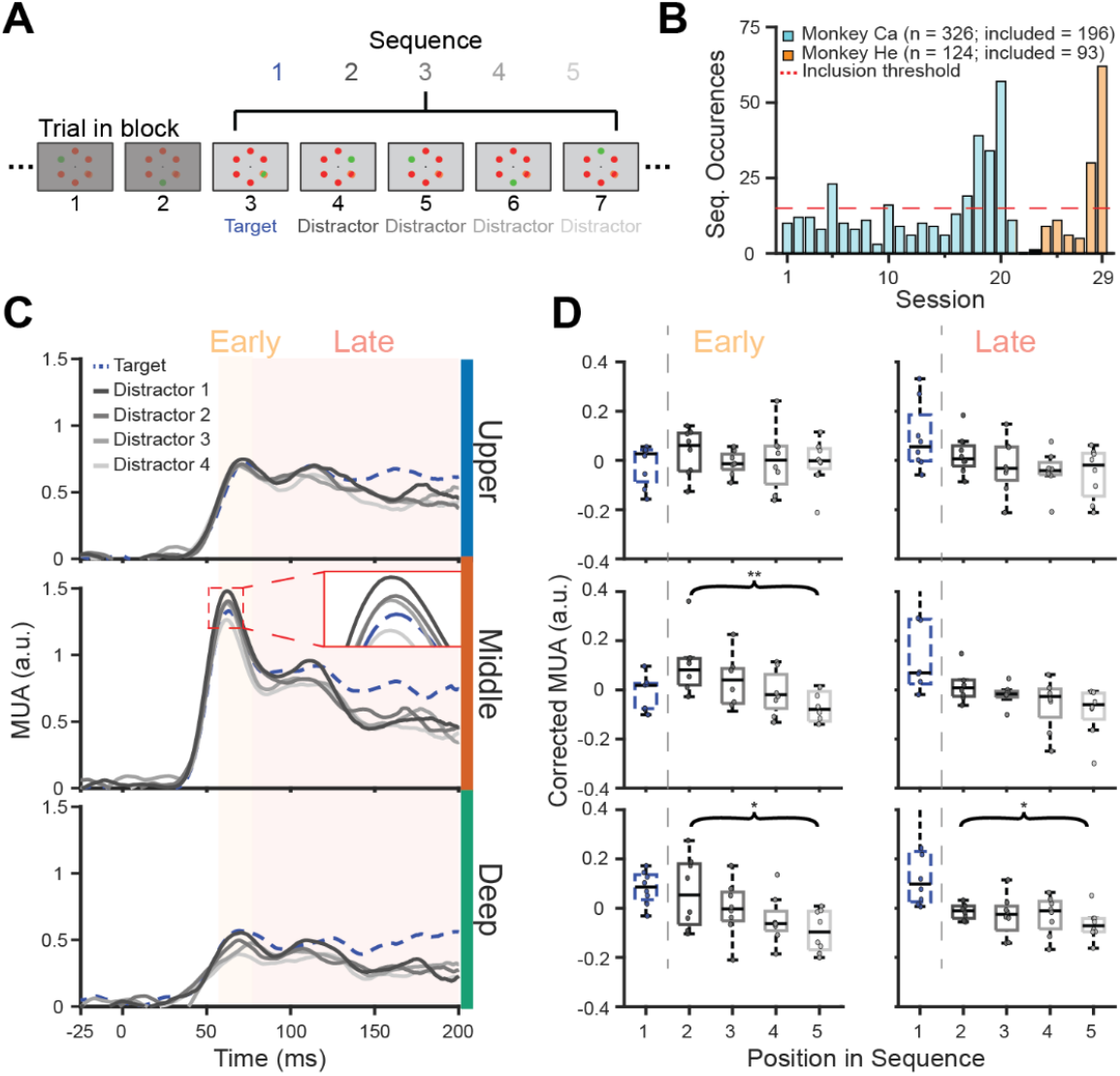
Feedforward neural adaptation contributes to improved distractor suppression. **A)** Schematic of the sequence of interest, consisting of a predictable target trial followed by four distractor trials, during which target and distractor features were held constant. **B)** Histogram of the number of qualifying sequences per session for each monkey; only sessions containing at least 15 such sequences were included, yielding 289 occurrences across 8 sessions (2 monkeys). **C)** Average multiunit response to each sequence position, shown separately for upper (top), middle (middle), and deep layers (bottom). Target responses are plotted as blue dotted lines and distractor responses in gray solid lines (dark to light with increasing sequence position); the inset in the middle panel shows responses between 50–75 ms for amplitudes between 1.2–1.5 a.u. **D)** Session-averaged, baseline-corrected responses (boxplots) for each sequence position and laminar compartment, quantified for an early visual response window (left; 58–78 ms after stimulus onset) and a late window (right; 78 ms after stimulus onset to 10 ms before saccade onset). The sequence makeup was a target followed by 4 distractors (T-D-D-D-D). For each session, the mean across all sequence positions was subtracted from each position. Monotonic decrease in distractor response was statistically tested using the Page’s L test (α = 0.05).

We statistically assessed this reduction by testing for monotonic trends in session-average responses across all distractor trials within each laminar compartment using Page’s L test (α = 0.05). Here, we distinguished between the early- and late-response periods (Figure 4D) to characterize potential differences in the feedforward response versus the period of recurrent activation. For the early response period, we averaged activity between 58 and 78 ms after stimulus onset (Figure 4D, left), consistent with the initial feedforward response period previously described [42]. At the level of the whole cortical column, we observed a significant monotonic reduction in the session averaged response across distractor repetitions (L = 225, p = 0.0011). When separated per laminar compartment, the median responses showed a significant monotonic reduction in activity in the middle (L = 221, p = 0.00510) and deep layers (L = 215, p = 0.0331). No significant effect was observed in the upper layers (L = 203, p = 0.357). For the late response period, we averaged the activity from 78 ms after stimulus onset up to 10 ms before saccade onset (Figure 4D, right). Consistent with the early-period findings, the whole cortical column likewise exhibited a significant monotonic reduction during this interval (L = 217, p = 0.0187). Here, we observed a significant reduction in the deep layers (L = 214, p = 0.0432), but not in the upper (L = 207, p = 0.196) or middle layers (L = 210, p = 0.110), although the middle layers were approaching significance and qualitatively reflect the adaptation profile. Together, these results indicate that the distractor suppression effect, at least to some degree, can be explained by neuronal adaptation to the more frequent distractor stimulus.

### More reliable target responses in the predictable context which predict behavior

While both TE and DS, and changes in TE and DS, are present across laminar compartments, it remains unclear whether their magnitude directly impacts behavioral outcomes in target identification. Previous work suggests that target, but not distractor responses in visual cortex predict RT [42]. However, this was not evaluated relative to stimulus predictability. To address this, we examined TE and DS stratified by RT. We analyzed response profiles separately in unpredictable and predictable contexts, rather than subtracting them, as this approach reveals more nuanced differences.

Separation by RT revealed a distinct pattern in target-evoked responses (Figure 5A). In the unpredictable context, average response profiles to target presentations differed significantly across behavioral outcome groups in all cortical layers, with the deep layers exhibiting the most robust effects (running k-test, p < 0.01; mean onset time = 141 ms). Importantly, differences across RT quartiles emerged during the feedforward visual sweep (∼50-60 ms following array onset) and persisted throughout the response period, indicating that the strength of the initial feedforward drive robustly predicted reaction time performance under unpredictable conditions [42].

**Figure 5.**
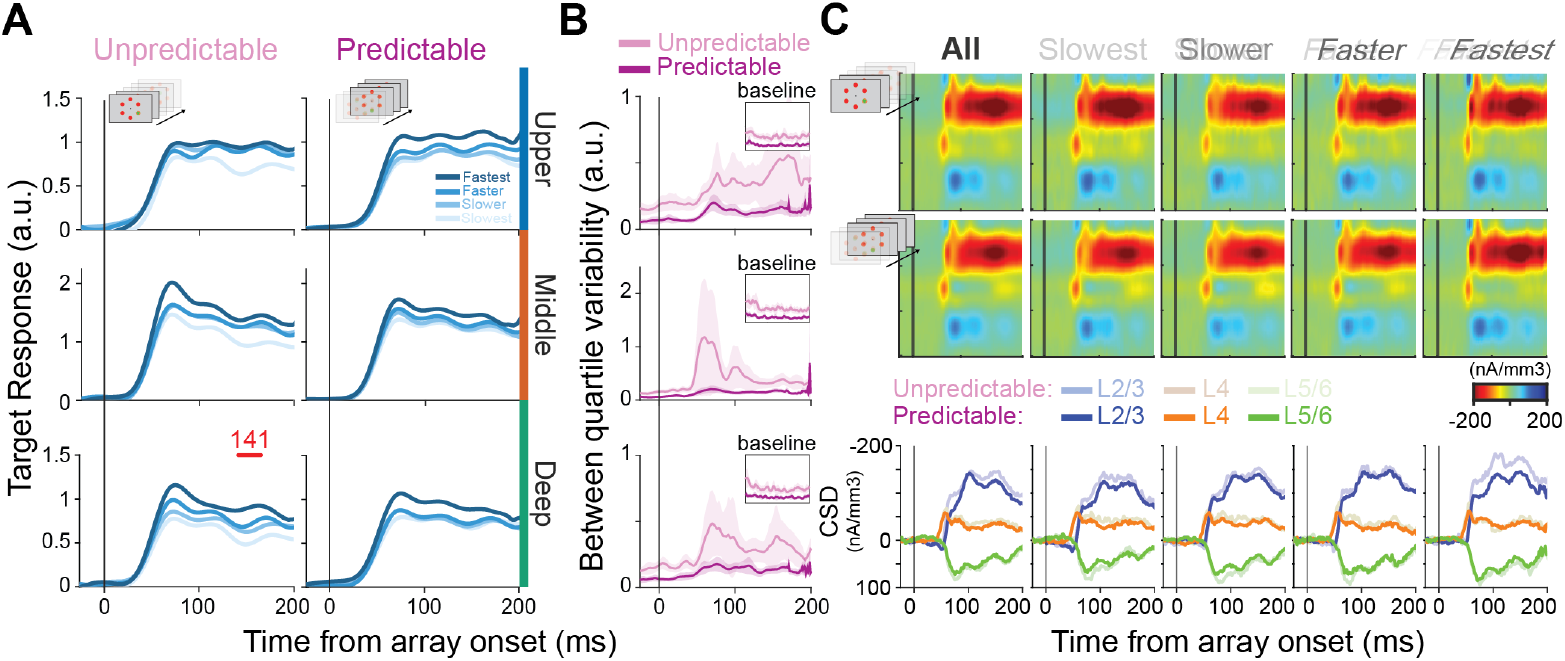
Useful attention capture through more reliable columnar sensory responses with target predictability. **A)** Average target response for unpredictable (left) and predictable (right) contexts, stratified by behavioral performance (reaction time quartile from slowest to fastest, color gradient). Significant differences between behavioral outcome groups are denoted by red bars atop each plot (running Kruskal-Wallis test; α = 0.01, minimum cluster length = 25 ms). **B)** Between quartile variability in both the unpredictable (pink) and predictable condition (purple), aligned to visual array onset. Inset displays the variability during the baseline period (-300 - 0 ms before visual array onset). **C)** CSD-plots for unpredictable (top), and predictable (middle) contexts, displayed for all trials (“All”) and stratified by behavioral performance (“Slowest” - “Fastest”). The bottom row displays the CSD response profiles per laminar compartment (superficial = blue; granular = orange; infragranular = green) with the predictable context in vibrant colors, and the unpredictable context in muted colors.

With predictability, target responses converged toward a more reliably strong and sustained activation profile (Figure 5A, right column). Slower trials exhibited relatively greater activity, whereas faster trials displayed responses similar or slightly reduced compared to the unpredictable condition, resulting in more consistent response profiles across reaction time outcome groups.

This is further illustrated by a marked reduction in between-quartile variability of target responses in the predictable context across all laminar compartments (Figure 5B). To be clear, variability here is quantified as the variance across the four RT quartile-average response profiles at each time point, where a higher value reflects greater divergence in neural activity between behavioral outcome groups. Interestingly, a reduction in variability from unpredictable to predictable contexts is already evident during the baseline period (Figure 5B inset), suggesting that baseline activity in unpredictable contexts may be more important in determining whether the correct stimulus captures attention. Baseline activity has been shown to impact the likelihood of attentional capture [42]. All in all, this suggests that predictability alters the cortical column by reducing neural response variability, thereby promoting more stable and reliable target response profiles and, consequently, more reliable, useful attentional capture across trials.

As a final step in assessing target-related changes, we investigated synaptic activity using CSD analyses (Figure 5C). CSD provides an indication of putative synaptic activation [70]. When examining the target responses in the unpredictable context, we observe an initial current sink in the middle layer (∼50 ms), followed by a pronounced sink in the upper layers (Figure 5C; “All” column). When quartiled by RT, we observe a pattern in which faster trials generally display stronger putative synaptic activation (Figure 5C, “Slowest” to “Fastest” columns), suggesting that faster target selection is associated with greater synaptic drive.

Surprisingly, we observe a reduction in putative synaptic activation with target predictability, most pronounced in the upper layers (Figure 5C). Note, this effect is only qualitative, and statistics only trend toward significance in the upper layers. This may suggest that while faster target selection is generally associated with stronger spiking responses to the target in predictable contexts, synaptic activation may conversely decrease.

### DISTRACTOR responses do not predict behavior

We next performed an identical analysis for the distractors (Figure 6). We observed that distractor responses, quartiled by RT, displayed overlapping response profiles in both the unpredictable and predictable contexts (Figure 6A). This suggests that, while DS is present, the strength of the distractor response is not predictive of behavior in either context. Corresponding CSD profiles across cortical layers displayed a similar pattern to that observed for target presentations: faster trials displayed more synaptic activity, while increased predictability led to a reduction in overall synaptic activity (Figure 6B). This further suggests that the changes in distractor suppression we observed likely result from neural adaptation rather than from active top-down modulation.

**Figure 6.**
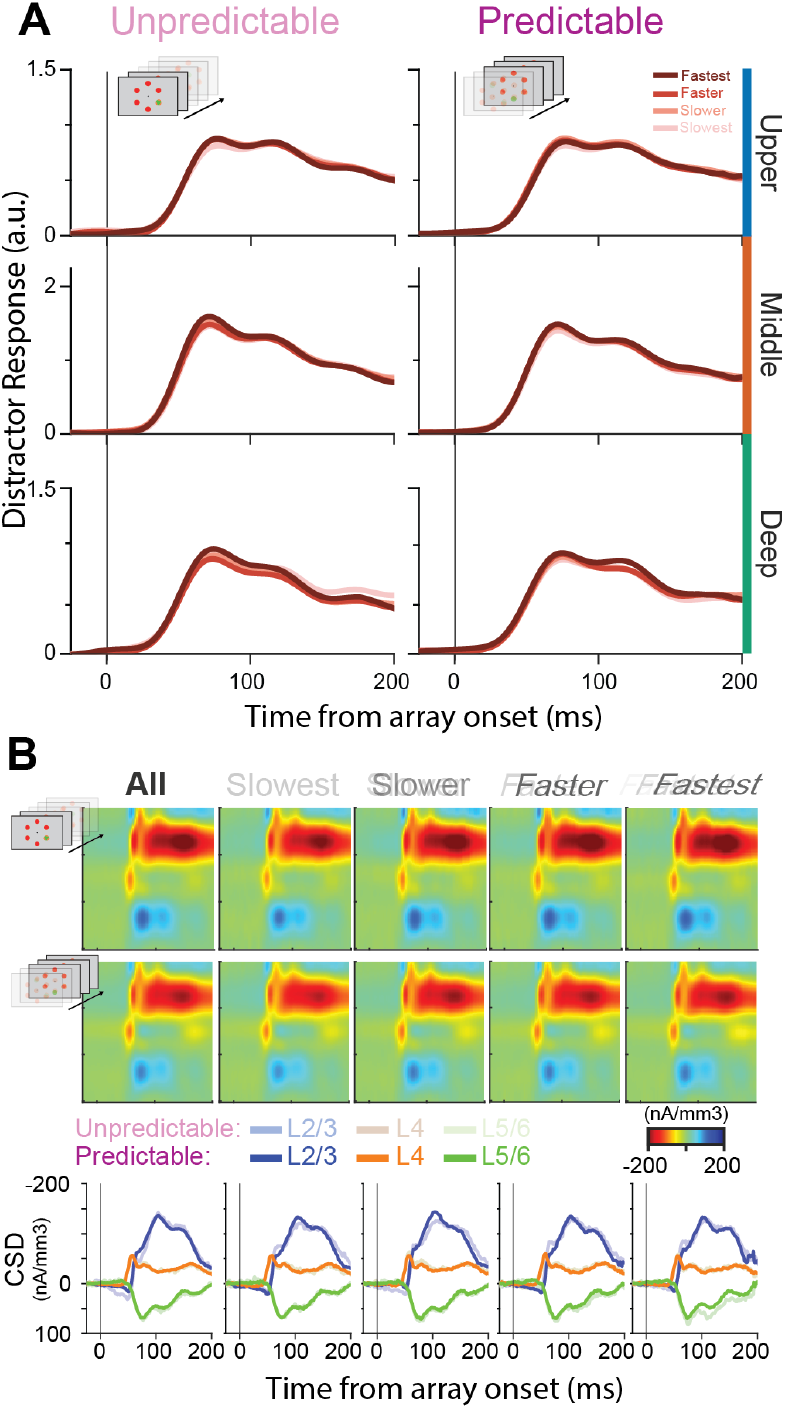
Distractor responses are not predictive of behavior in either predictability context. **A)** Average distractor responses for unpredictable (left) and predictable (right) contexts, stratified by behavioral performance (reaction time quartile from slowest to fastest, color gradient). Significant differences between behavioral outcome groups are denoted by red bars atop each plot (running Kruskal-Wallis test; α = 0.01, minimum cluster length = 25 ms). **B)** CSD plots for unpredictable (top), and predictable (middle) contexts, displayed for all trials (“All”) and stratified by behavioral performance (“Slowest” - “Fastest”). The bottom row displays the CSD response profiles per laminar compartment (superficial = blue; granular = orange; infragranular = green) with the predictable context in vibrant colors, and the unpredictable context in muted colors

## Discussion

Predictable attentional capture through priming in pop-out visual search facilitates faster and more accurate attentional selection [48,57]. However, how predictability modulates the responses of the underlying neural circuitry that supports this improvement remained an open question. Here, we examined how predictability modulates attentional selection and sensory processing across cortical layers in area V4. By characterizing laminar and temporal response dynamics, we revealed how the predictability of attentional capture reorganizes cortical processing to facilitate attentional selection primarily through changes in feedforward signaling.

### Predictability fosters more reliable feedforward target discrimination

Manipulating target predictability induced a seemingly marked reconfiguration of feedforward processing within V4. In the unpredictable sensory context, target selection dynamics exhibited pronounced laminar differences when related to behavior (Figure 3A). The fastest RT trials were characterized by early and strong feedforward activation in the middle and deep layers, followed by recruitment of the upper layers around the time when feedback to V4 is typically observed (>100 ms post-stimulus onset). In slower trials, this laminar cascade of selection was delayed and weakened, suggesting that successful performance in unpredictable contexts hinges on the strength and timing of the initial feedforward drive. Moreover, the slowest (and “slower”) trials also showed an erroneous early selection (i.e., selection profiles below 0), indicating that the feedforward drive may have selected a distractor stimulus before course-correcting around the time of putative feedback, likely due to recurrent modulation.

Once the sensory context became predictable, target selection became virtually simultaneous across laminar compartments, with diminished differentiation between behavioral outcome groups. Additionally, early erroneous selections were not observed at the population level, unlike in the unpredictable condition. These changes indicate a more reliable feedforward processing mode, which could yield more reliable behavioral responses without *necessitating* top-down enhancement of relevant target representations. However, it is worth noting that target selection profiles become more pronounced around the time of feedback to V4, when spatial selective attention is typically deployed.

The early onset of these modulations argues against a dominant role for rapid reactive top-down feedback from prefrontal or parietal sources in producing the changes in the strength and speed of attentional enhancement in V4 in this paradigm. For instance, an alternative hypothesis would suggest that a downstream brain area discriminates the target stimulus more rapidly from sensory information that does not differ across contexts, and that feedback from that area to V4 is more rapidly deployed to produce the observed changes in target enhancement and distractor suppression [46]. This would manifest as changes in selection profiles around the time of putative feedback, but not before. Instead, our findings suggest that feedforward signaling is already altered in this predictable context, supporting behavioral facilitation. These results support a model in which local sensory circuits contribute significantly to attentional selection rather than merely serving as static recipients of top-down modulation [42,71,72].

### More potent distractor suppression as a bottom-up neural response adaptation

While behavior seemed better predicted by the neural response to the target stimulus in either context, the changes in distractor processing likely contributed to the target stimulus’s enhanced discriminability. After all, a lower distractor response would create a greater “target-signal” in “distractor-noise” in a signal detection theory framework [73]. Here, we examined the source of the changes in distractor suppression between unpredictable and predictable search contexts. While active distractor suppression via putative selective feedback has been observed in area V4 [68], presumably via attentional control structures in the prefrontal cortex [74], our results do not indicate that this is the only mechanism employed here. The onset latency of changes in distractor suppression and its progression from middle to upper and deep layers are suggestive of a change in feedforward processing of the distractor stimulus (Figure 3B). The most parsimonious explanation for this change is neural adaptation to the distractor stimulus [65,69,75–78]. This is likely in this paradigm, given that five out of six stimulus locations contain a distractor on a given trial. Consequently, for any neuron’s RF that overlaps with a fixed stimulus location, the probability of repeated distractor presentation across consecutive trials is high. Such repeated exposure to the same distractor feature may drive stimulus-specific adaptation, progressively attenuating the neuron’s response to that stimulus over time [65,69,75–81]. Our results align with this hypothesis, showing a reduction in the initial feedforward response when the same distractor stimulus was presented across consecutive trials. Therefore, the distractor suppression observed in our context likely results from neural adaptation rather than active top-down suppression. This is consistent with so-called “BELIEF” models of predictive processing, which use feedforward neural adaptation to explain diminished responses to expected stimuli (e.g., more frequent stimuli) [82].

### Downstream implications of feedforward adaptation

A key question raised by our findings is how the experience-dependent changes in feedforward processing we observe in V4 propagate through the visual hierarchy to influence behavior. Work in the frontal eye fields (FEF) provides a useful window into this question. Under normal conditions, FEF neurons, like most prefrontal neurons, do not exhibit feature selectivity. However, in monkeys trained exclusively on targets of a given feature (e.g., color), FEF neurons develop selectivity for that feature, occurring soon after array presentation and independent of location within the visual field [83].

This feature selectivity could result from inherited upstream changes in V4, such as what we observe in this study. The anatomical projection from V4 to FEF is well established [84–87]. Long-term adaptation to the same distractor feature could in turn produce a selectivity for the target feature. In other words, the diminished responses in sensory cortical areas could result in less excitation of higher-order neurons (e.g., in FEF) under the distractor stimulus conditions thereby producing a selectivity for the target feature which is less frequent and less adapted. This poses an interesting hypothesis for future work to establish the origins for spontaneous feature selectivity in prefrontal cortex.

### Proactive feedback vs. local functional reorganization for enhanced target signaling

While the changes in distractor suppression are most parsimoniously explained by neural adaptation, the source of changes to target processing is less clear. As distractor responses did not vary with behavior, nor were there notable laminar differences in distractor responses in either predictability context, the observed changes in feedforward selection for behavior likely stem primarily from changes in target processing. Previous work suggests that prestimulus changes in cortical columnar processing may promote representations of relevant features for attentional capture (e.g., enhanced baseline activity in red-preferring cortical columns during predictable red-stimulus search) [42]. Furthermore, the findings here suggest reduced trial-to-trial variability in neural responses (and baseline activity) in predictable contexts, as indicated by observing less variable target responses across RT quartiles (Figure 5B). However, how exactly predictability establishes more homogenous feedforward selection profiles across the layers of V4 remains unclear. One obvious question is whether the changes we observe stem from bottom-up-driven changes in feedforward processing or from proactive top-down modulation affecting the target processing in predictable sensory contexts.

### More efficient feedforward attentional selection

The transition to a predictable sensory context was accompanied by a putative, albeit only qualitative, reduction in synaptic activity, particularly in the upper cortical layers when a target was present in the RF (Figure 5C). Remarkably, this was the case despite concomitant increases in spiking activity in the same condition. While only a modest effect size was seen here, the observation is notable given the considerable interest in predictive processing as an optimization of cortical information passing. This suggests that cortical microcircuits adopt a more efficient, streamlined mode of processing in predictable contexts. How exactly putative reductions in synaptic activation yield greater neural spiking responses remains unclear from the analyses performed here. One possible explanation is that the predictable context is associated with changes in the spike timing of brain areas that provide input to V4, such that stronger downstream activity can be facilitated by fewer, more synchronous spikes [9,15,35,36]. Further investigation into the role of spike timing in predictable attentional capture remains to resolve this curious observation.

## Conclusions

Behavior is facilitated in predictable contexts. The findings here have implications for models of how predictive processing is instantiated in the visual cortex to support this behavioral facilitation. Feedforward processing of predicted relevant stimuli becomes stronger and more reliable, and processing resources dedicated to irrelevant stimuli are reduced, likely via automatic bottom-up adaptations. These findings suggest a critical role for changes in feedforward processing for facilitating attentive behavior in predictable contexts.

## Data and code availability

All data used in this study are openly available on Zenodo (https://doi.org/10.5281/zenodo.19512221). All code used to generate the results in this study from the dataset is available on GitHub (https://github.com/JiskGroot/GrootWesterberg2026).

## Acknowledgments

This work was supported by the NIH National Eye Institute [grant numbers: R01EY019882, R01EY008890, P30EY008126], the Dutch Research Council (NWO) [grant number: VI.Veni.232.110], and the Natural Sciences and Engineering Research Council of Canada [grant number: RGPIN-2022-04592]. J.A.W. was supported by fellowships from the NIH National Eye Institute [grant numbers: F31EY031293, T32EY007135] and the International Human Frontier Science Program Organization [grant number: LT0001/2023-L]. Imaging support was provided by the Vanderbilt University Institute for Imaging Science through a grant from the National Institutes of Health, Office of the Director [grant number: S10OD021771].

## Author contributions

Conceptualization, J.D.S., J.A.W.; Data Collection, J.A.W.; Formal Analysis, J.J.G., J.A.W.; Data Visualization, J.J.G.; Original Draft, J.J.G., J.A.W.; Revisions and Final Draft, J.J.G., J.D.S., J.A.W..

## Methods

### Animal care

Two male macaque monkeys (*Macaca radiata*; monkey Ca, He) participated in this study. All procedures were in accordance with the National Institutes of Health Guidelines and the Association for Assessment and Accreditation of Laboratory Animal Care International’s Guide for the Care and Use of Laboratory Animals and approved by the Vanderbilt Institutional Animal Care and Use Committee in accordance with United States Department of Agriculture and United States Public Health Service policies. Animals were pair-housed. Animals were on a 12-hour light-dark cycle, and all experimental procedures were conducted in the daytime. Each monkey received nutrient-rich, primate-specific food pellets twice a day. Fresh produce and other forms of environmental enrichment were given at least five times a week.

### Surgical procedures

All surgical procedures were performed under aseptic conditions. Anesthesia was conducted with animals under N_2_O/O_2_, isoflurane (1-5%) anesthesia mixture. Vital signs were monitored continuously. Expired PCO_2_ was maintained at 4%. Postoperative antibiotics and analgesics were administered while animals remained closely observed by veterinarians and staff. Monkeys were implanted with a custom-designed head post and MR-compatible recording chamber using ceramic screws and biocompatible acrylic. A craniotomy over V4 was opened, and a recording chamber was placed above the craniotomy.

### MRI

MR images were taken from anesthetized animals placed inside a 3T MRI scanner (Philips) to localize chambers and guide linear electrode penetrations perpendicular to the cortical surface. T1-weighted 3-dimensional MPRAGE scans were acquired with a 32-channel head coil for SENSE imaging. Images were acquired using a 0.5 mm isotropic voxel resolution with the following parameters: repetition time 5 s, echo time 2.5 ms, and flip angle 7°.

### V4 identification

Recordings took place on the convexity of the prelunate gyrus in approximately the dorsolateral, rostral aspect of the V4 complex, where receptive fields are located perifoveal, at about 2–10 degrees of visual angle (dva) eccentricity in the lower contralateral visual hemifield [88]. Laminar recordings occurred where the array could be positioned orthogonal to the cortical surface, as verified by MRI and neurophysiological criteria (i.e., overlapping receptive fields, CSD). Recording sites were also confirmed via histological staining by dipping the electrode arrays in diiodine before the final recordings in monkey He [48].

### Priming of pop-out task

Monkeys viewed arrays of stimuli on a CRT monitor with a 60 Hz refresh rate at a 57 cm distance. Stimulus presentations and task timing were controlled using TEMPO (Reflective Computing). Visual presentations were monitored with a photodiode positioned on the CRT monitor to align offline electrophysiological signals. Red (CIE coordinates: x=0.648, y=0.331) and green circles (CIE coordinates: x=0.321, y=0.598) were used as stimuli, rendered isoluminant to a human observer at 2.8 cd/m^2^ on a uniform gray background. As we are limited to two colors and cannot account for potential differences in perceived brightness between macaques, we qualify our two stimuli as distinct ‘features’ at the intersection of color and luminance information. Nonetheless, we report the colors used in this study for the ideal human observer. Cone excitation was computed from the CIE coordinates and luminance [89]. The following cone excitations were measured for red: ε_*L*_=2.37, ε_*M*_=0.43, ε_*S*_=0.0014; green: ε_*L*_=1.74, ε_*M*_=1.06, ε_*S*_=0.0030; and the background: ε_*L*_=1.86, ε_*M*_=0.94, ε_*S*_=0.023. Cone contrasts for red stimulus were: *C*_*L*_=0.27, *C*_*M*_=-0.54, *C*_*S*_=-0.94; and the green stimulus: *C*_*L*_=-0.06, *C*_*M*_=0.13, *C*_*S*_=-0.87.

Trials were initiated when monkeys fixated within 0.5 dva of a small, white fixation dot (diameter = 0.3 dva). The time between fixation acquisition and array presentation varied between 750 and 1250 ms, taken from a nonaging foreperiod function to eliminate any potential temporal effect of stimulus expectation [90–92]. Following the fixation period, the stimulus array comprising six items was presented. Stimuli were scaled with eccentricity at 0.3 dva per 1 dva eccentricity, so they were smaller than the estimated V4 receptive field size (0.84 dva per 1 dva eccentricity [93]). The polar angle positioning of the items relative to fixation varied from session to session, so one item of the stimulus array was positioned at the center of the population receptive field under study. Items were spaced such that only one item was in the V4 receptive field, with uniform spacing in polar angle and equal eccentricity.

Monkeys engaged in a search task while viewing the stimulus array. One item in the array was a different feature (red or green, respectively) from the others. The position of the oddball on each trial was randomly chosen with equal probability for any of the positions (16.6%). Monkeys earned fluid reward for shifting their gaze directly to the oddball item within 1000 ms of array presentation and maintaining fixation within a 2–5 dva window around the oddball for 500 ms. Eye movements were monitored continuously at 1 kHz using an infrared corneal reflection system (SR Research). If the monkey failed to look at the oddball, no reward was given, and a 1-5 s timeout ensued. Trials were organized into blocks, and the animal searched for the same target feature for typically 5-15 repetitions. The target feature remained the same, but the target location varied randomly. Completing the block resulted in the target and distractor features swapping.

### Neurophysiology

Laminar extracellular voltages were acquired at 24.4 kHz resolution using a 128-channel PZ5 Neurodigitizer and RZ2 Bioamp processor (Tucker-Davis). Raw signals were output between 0.1 Hz and 12 kHz. Data were collected from 2 monkeys (left hemisphere, monkey Ca; right hemisphere, He) across 70 recording sessions (n=31, monkey Ca; n=39, He) using 32-channel linear microelectrode arrays with 0.1 mm interelectrode spacing (Plexon). During each recording session, electrode arrays were introduced into the prelunate gyrus through the intact dura mater using a custom micromanipulator (Narishige). Electrode arrays were positioned to span all layers of V4 and had a subset of electrodes positioned outside the cortex. 29 (n=21, monkey Ca; n=8, He) of 70 sessions were included in the final analysis. The remaining 41 sessions were excluded from the analysis because they lacked a discernible CSD profile for laminar alignment, were not orthogonal to the cortical surface, or had insufficient priming blocks.

### Receptive field mapping

Monkeys performed a receptive field mapping task before the main task to determine the orientation and eccentricity of the visual receptive fields. Monkeys fixated for 400–7000 ms while a series of 1–7 stimuli spanned the visual field contralateral to the recording chamber. Stimuli were five high-contrast concentric white and black circles that scaled in size with eccentricity (0.3 dva per 1 dva eccentricity). In all recording sessions, stimuli could appear in a random location. These random locations spanned the lower visual quadrant contralateral to the recording chamber. Location spacing was in 5° angular increments relative to fixation and eccentricities ranging from 2 dva to 10 dva in 1 dva increments. Each stimulus was presented for 200–500 ms with an interstimulus interval of 200–500 ms. If the animal maintained fixation during the stimulus presentation sequence, they received a juice reward. This receptive field mapping task measured multiunit activity and gamma power (30-90 Hz) as well as evoked local field potentials (LFPs, 1–100 Hz) across all recording sites. Online, we measured the response across visual space for each recording site. Recordings proceeded to the feature search task if there was qualitative homogeneity of receptive fields along the depth. Receptive field overlap for these data has been reported previously [21]. The receptive field center was chosen as the stimulus location that evoked the largest response along the depth of the recording sites. Following receptive field identification, the stimulus array in the feature search task was then oriented so that its eccentricity coincided with the location of the receptive field (eccentricity: 3–10 dva), and a single array item was placed at the center of the receptive field (size: 0.9–3 dva).

### Laminar positioning

Positions of the individual recording sites relative to the layers of V4 were determined using current source density (CSD) analysis. CSD reflects putative local synaptic currents (net depolarization) resulting from excitatory and inhibitory postsynaptic potentials [70]. CSD was computed from the raw neurophysiological signal by taking the second spatial derivative along the electrode contacts [62,63,65,94,95]. CSD activation following the presentation of a visual stimulus reliably produces a specific pattern of activation, which can be observed in the primate visual cortex [62,63], including V4 [18,21,96–98]. Specifically, current sinks following visual stimulation first appear in the granular input layers of the cortex and then ascend and descend to the extragranular compartments. To compute the CSD from the LFP, we used the previously described procedure [95]:

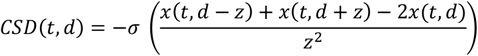

The CSD at timepoint t and cortical depth d is the sum of voltages x at electrodes immediately above and below (z is the interelectrode distance) minus 2 times the voltage at d divided by the interelectrode-distance-squared. That computation yields the voltage local to d. To transform the voltage to current, we multiplied that by -σ, where σ is a previously reported estimate of the conductivity of the cortex [99]. For each recording session, we computed the CSD and identified the initial granular layer (L4) input sink following visual stimulation. Sessions were aligned using the bottom of the initial feedforward input sink as a functional marker. We defined the size of individual laminar compartments uniformly relative to space. Throughout, ‘middle’ refers to the estimate of the granular input layer 4 (0.5 mm space above the CSD initial sink functional marker), ‘upper’ refers to the estimate of supragranular layers 2 and 3 (0.5 mm space above the L4 compartment), and ‘deep’ refers to the estimate of infragranular layers 5 and 6 (0.5 mm space below the L4 compartment).

### Multiunit envelope

Spiking activity at the level of multiunits was used for control analyses as it reliably reflects the neural population dynamics [100]. Detection of multiunit activity was achieved through previously described means [101]. This method has proved useful across brain areas and research groups [48,64,102–107]. Briefly, broadband neural activation was filtered between 0.5 and 5 kHz, the predominant range of spiking activity. The signal was then full-wave rectified and filtered again at half the original high-pass filter (0.25 kHz), thereby estimating the power of the multiunit activity. For filtering, we used a 4^th^-order Butterworth filter.

### Data analysis

#### Preprocessing

Analyses were restricted to trials that were correctly performed. Neural signals were aligned to stimulus onset and normalized at the single-trial level using a z-score approach. Each data point was divided by the standard deviation of the 250-ms pre-stimulus baseline period. Baseline correction was then applied by subtracting the mean baseline activity from each trial. To minimize artifacts arising from saccadic eye movements, data from 10 ms before saccade onset until the end of the trial were excluded, except for saccade-aligned analyses. Finally, a 15-ms bidirectional moving average filter was applied to smooth the data.

Data were classified into unpredictable and predictable contexts based on trial number relative to a feature change: the first two trials post-feature change were considered unpredictable, while trials three through fifteen were designated as predictable.

#### Reaction time stratification

Within each session, trials were sorted by reaction time and divided into quartiles [42]. This stratification generated four behavioral groups per priming condition (fastest, fast, slower, slowest), enabling comparisons of neural activity as a function of response speed. Only correct trials were considered in the binning process.

#### Target selection time, target enhancement, and distractor suppression

Target selection time (TST) was defined as the earliest time point post-array onset at which population activity in V4 significantly differentiated the target from distractor trials. TST was estimated by comparing MUA averaged across sessions at the channel level and identifying the first time point when target-related activity exceeded distractor-related activity significantly.

To further elucidate mechanisms underpinning target selection, we quantified target enhancement (TE) and distractor suppression (DS) as changes in neural responses to target and distractor conditions, respectively, comparing predictable to unpredictable states using running rank-sum tests.

#### Neural adaptation

To test whether distractor suppression reflected neural adaptation to repeated distractor presentations, we identified trial sequences in which the same distractor stimulus was presented consecutively following a predictive target. Sequences were required to start with a target trial during the predictable context, followed by at least four consecutive distractor trials within the same target-feature block, such that target and distractor features remained constant across the entire sequence. We only included sessions that contained at least 15 such sequences. For the analysis we distinguished between the early response phase (58–78 ms after stimulus onset), and late response phase (78 ms after stimulus onset to 10 ms before saccade onset).

#### Statistical analysis

Statistical comparisons of behavioral performance were performed using a one-way repeated measures ANOVA with Trial position as the within-subject factor (15 levels; α = 0.05), followed by post-hoc paired t-tests between consecutive trial positions (two-sided, α = 0.05), based on session-wise averages pooled across animals (n = 29). Reaction time differences between unpredictable and predictable contexts were assessed using the Wilcoxon signed-rank test (two-sided; α = 0.05), conducted separately per animal (Monkey Ca: n = 21 sessions; Monkey He: n = 8 sessions). Neural adaptation was assessed using Page’s L test (one-sided, decreasing trend; α = 0.05) [108] on session-demeaned averages across four consecutive distractor trials (D1-D4; k =4), restricted to sessions with ≥15 complete sequences (n = 8). Comparisons of MUA between unpredictable and predictable contexts were conducted using a running Wilcoxon rank-sum test against a pre-stimulus baseline distribution (-250 to 0 ms; two-sided; α = 0.01, minimum cluster length = 25 ms) based on session- and channel-wise averages (n = 145 per compartment; 29 sessions × 5 channels). Per-channel target versus distractor differences were assessed with a running Wilcoxon rank-sum test (two-sided; α = 0.01, minimum cluster length = 25 ms) applied to pooled individual trials across sessions. Differences between reaction time groups were evaluated using a running Kruskal-Wallis test (α = 0.01, minimum cluster length = 25 ms) on session- and channel-wise averages (n = 145 per compartment). All error bars and shaded regions represent the mean ± 95% CI (±1.96 × SEM).

